# Boosting innate immunity: *Asaia* bacteria expressing a protein from *Wolbachia* determine macrophage activation and killing of *Leishmania*

**DOI:** 10.1101/2020.06.22.164145

**Authors:** Ilaria Varotto-Boccazzi, Sara Epis, Irene Arnoldi, Yolanda Corbett, Paolo Gabrieli, Moira Paroni, Riccardo Nodari, Nicoletta Basilico, Luciano Sacchi, Marina Gramiccia, Luigi Gradoni, Vito Tranquillo, Claudio Bandi

## Abstract

Leishmaniases are severe vector-borne diseases affecting humans and animals, caused by *Leishmania* protozoans. Immune polarization plays a major role in determining the outcome of *Leishmania* infections: hosts displaying M1-polarized macrophages are protected, while those biased on the M2 side acquire a chronic infection, that could develop into an overt and potentially deadly disease. The identification of the factors involved in M1 polarization is essential for the design of therapeutic and prophylactic interventions, including vaccines. Infection by the filarial nematode *Dirofilaria immitis* could be one of the factors that interfere with leishmaniasis in dogs. Indeed, filarial nematodes induce a partial skew of the immune response towards M1, likely caused by their bacterial endosymbionts, *Wolbachia*. Here we have examined the potential of *Asaia*^WSP^, a bacterium engineered for the expression of the *Wolbachia* surface protein (WSP), as an inductor of M1 macrophage activation and *Leishmania* killing. Macrophages stimulated with *Asaia*^WSP^ displayed a strong leishmanicidal activity, comparable to that determined by the choice-drug amphotericin B. Additionally, *Asaia*^WSP^ determined the expression of markers of classical macrophage activation, including M1 cytokines, ROS and NO, and an increase in phagocytosis activity. *Asaia* not expressing WSP also induced macrophage activation, although at a lower extent compared to *Asaia*^WSP^. In summary, our study, while providing a strong evidence for the immune-stimulating properties of *Wolbachia*, highlights the translational potential of *Asaia*^WSP^ in the areas of the immune-prophylaxis and therapy of leishmaniases, as well as of other diseases that could be subverted by M1 macrophage activation.

## Introduction

Naïve macrophages (M0) can differentiate into two major, functionally distinct, subtypes: the classically activated- and the alternatively activated-macrophages (indicated as M1 and M2, respectively). These myeloid cells play crucial roles non only in the immunity towards microbial and parasitic infections, but also in wound healing, tissue repair and in cancer progression or regression (Zhu et al., 2015; Ruytinx et al., 2018). Classically activated macrophages are the pro-inflammatory subtype with microbicidal properties, and the activation of the M1 response is intrinsically associated with increased phagocyte activity and killing of intracellular pathogens, through the production of reactive oxygen species (ROS) and nitric oxide (NO) (Atri et al., 2018); M1 macrophages are also crucial in anti-cancer immunity. The M2 phenotype is an anti-inflammatory/regulatory subtype, that plays a role in the resolution of inflammation and in tissue repair, as well as in tumor progression, and in variety of diseases associated with excessive antibody production (Weagel et al., 2015). Macrophage activations on the M1 or M2 side are associated with corresponding pathways in the polarization of T-helper lymphocyte cells, known as Th1 or Th2 (Parisi et al., 2018).

A parasitic infection that is paradigmatic in terms of its clinical outcome in relation with the M1 or M2 polarization is leishmaniasis. The general consensus is that during a *Leishmania* infection the development of a M1/Th1 response is associated with the production of proinflammatory cytokines such as TNFα, IL-12, and IFNγ and the release of ROS and NO, with the killing of the parasites, therefore with a protective immunity. On the other hand, a M2 response is associated with anti-inflammatory cytokine production, such as IL-4/IL-13, IL-10, TGFβ, M-CSF, expression of *arginase I* (with reduced NO production), inhibition of inflammation, parasite survival, and thus disease progression. In summary, while M1 activation is crucial for a successful elimination of *Leishmania* parasites, the M2 polarization, besides being unprotective, is in general associated with disease severity, also in relation with immune-complex pathology (Rossi & Fasel 2018).

Therefore, one of the major aims in leishmaniasis research is the identification of molecules with immunotherapeutic properties, i.e. molecules capable of modulating the immune response, to be used alone, in combination with drugs, or as vaccine adjuvants. Immunotherapy is already proposed for the control of several diseases, e.g. cancer, allergies, and viral infections, including COVID-19 (Roatt et al., 2014; Naran et al., 2018; Corey et al., 2020). In visceral leishmaniasis, patients non-responding to conventional chemotherapy have been treated with success through combination therapies with various immunomodulators, e.g. MDP13, IFNγ, IL-12, and the bacterium Bacille Calmette-Guérin (BCG) (El-On, 2009; Roatt et al., 2014). In addition, BCG has also been used in cutaneous leishmaniasis, in combination with a lysate of *Leishmania* parasites, with positive therapeutic effects (Mayrink et al., 1992; Convit et al., 2003).

In this study, we suggest that bacteria of the genus *Wolbachia* represent a promising source of molecules capable of stimulating and modulating innate immunity, with the potential to be exploited in immunotherapy and prophylaxis. In insects, *Wolbachia* has been shown to be a potent activator of innate immunity, able to determine the upregulation of several immune effectors such as antimicrobial peptides, autophagy-related proteins, and ROS (Zug & Hammerstein, 2015; Epis et al., 2020). Indeed, the successful use of *Wolbachia* to block the transmission of viruses by mosquitoes has in part been associated with this immune-activating capacity (Moreira et al., 2009; Rancès et al., 2012; LePage & Bordenstein, 2013). On the other hand, *Wolbachia* from filarial nematodes (or its surface protein, WSP) has been shown to activate macrophages through the stimulation of innate-immunity receptors, determining a M1/Th1-type activation (Saint André et al., 2002; Brattig et al., 2004). In summary, there is strong evidence that *Wolbachia* is an effective inductor of innate immunity, in insects and in mammals, and its surface protein WSP represents a promising candidate immunomodulator, with pro-M1 properties.

The exploitation of *Wolbachia* in macrophage stimulation experiments is however hampered by the characteristics of this bacterium: it is an obligate intracellular symbiont and it is not culturable in cell-free media, and thus not easy to be used in controlled *in vitro* and *in vivo* experiments. A strategy to deliver immunomodulators to hosts, for therapeutic or prophylactic purposes, is to engineer culturable non-pathogenic bacteria for their expression; the engineered bacteria are then administered to the host, through different routes (Berlec et al., 2019). For example, Jacouton and colleagues (2019) have modified a *Lactococcus lactis* strain for the expression of the cytokine IL-17A, with tumor prevention in a mouse model, after intranasal delivery of the engineered bacterium. In this context, we recently selected an acetic acid bacterium, *Asaia* sp., as a bacterial vehicle for the expression of the *Wolbachia* surface protein, generating the chimeric symbiont *Asaia*^WSP^ (Epis et al., 2020). Our hypothesis is that *Asaia*^WSP^ should determine innate immune activation with a M1 bias, thus conferring protection against M1-impaired infections. In order to test the potential of *Asaia*^WSP^ as an immunomodulating agent, we assayed its capability to stimulate the killing activity of macrophages towards *Leishmania infantum,* using a macrophage cell line, and determining the pattern of immune-activation of *Asaia*-stimulated cells (Fig. 1).

**Fig. 1.**
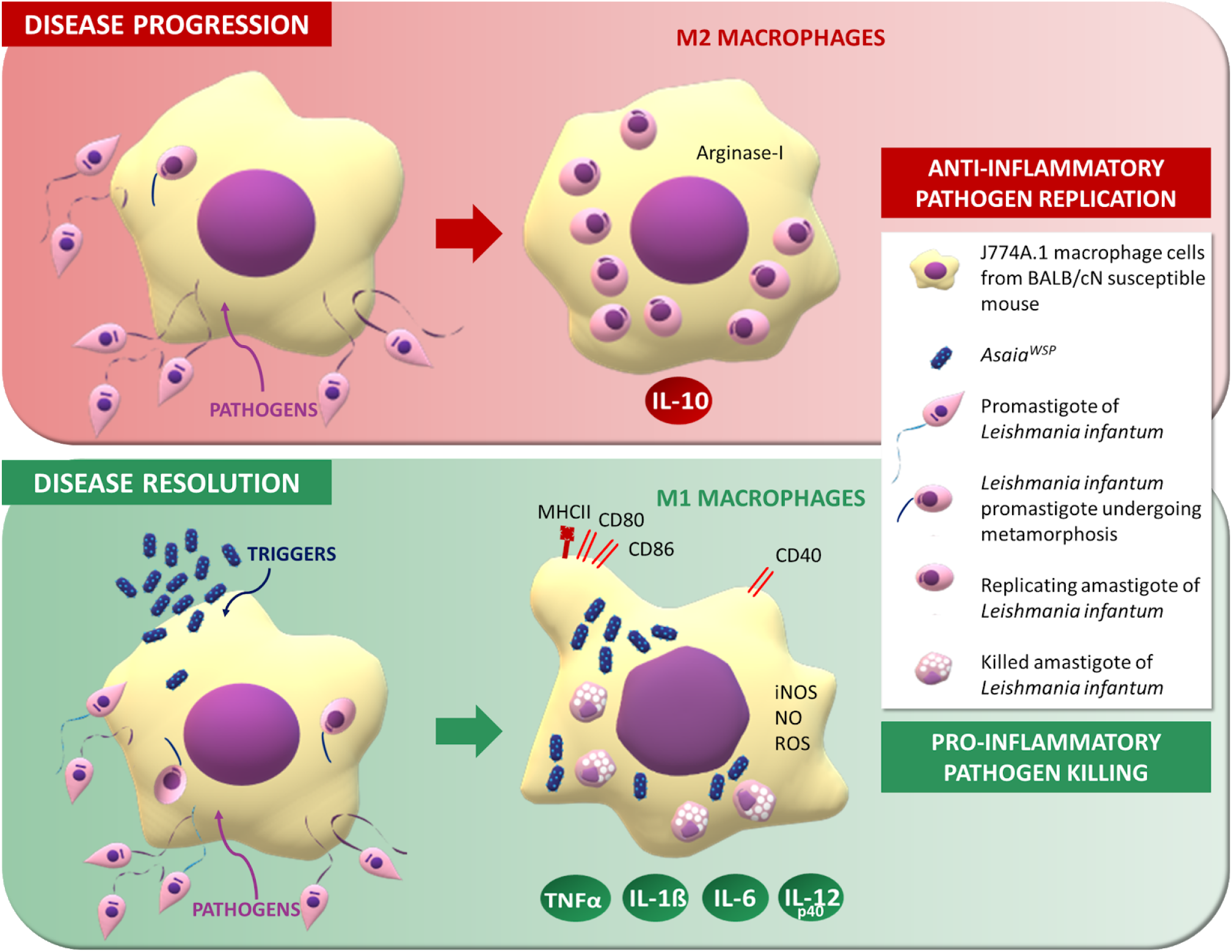
Graphical representation of the proposed mechanism of macrophage activation. The trigger *Asaia*^WSP^ acts as a polarizing agent, stimulating phagocytosis and inducing the release of pro-inflammatory mediators and microbicidal molecules. *Asaia*^WSP^ determines an anti-leishmanial effect with a reduction of the number of intracellular parasites. TNFα: tumor necrosis factor α; IL: interleukin; ROS: reactive oxygen species; iNOS: inducible nitric oxide synthase; NO: nitric oxide.

## Results

We planned our experiments starting from the engineered strains *Asaia*^pHM4^ and *Asaia*^WSP^; these had been derived from the wild-type strain *Asaia* SF2.1, using the plasmid vector pHM4, either “empty” (thus generating *Asaia*^pHM4^) or containing the WSP cassette (thus generating *Asaia*^WSP^) (Epis et al., 2020). The first set of experiments aimed at assessing whether incubation with *Asaia*^pHM4^ or *Asaia*^WSP^ stimulates the microbicidal activity of macrophages, on bacteria (*Asaia* themselves and *Staphylococcus epidermidis*) and on *Leishmania infantum* (hereafter *Leishmania*). Secondly, we characterized the pattern of macrophage activation determined by *Asaia*, focusing on markers of M1 polarization.

### Uptake and survival of bacteria in macrophages

The microbicidal activity of macrophages on bacteria was determined through the analysis of two processes: i) phagocytosis *sensu stricto*, i.e. the internalization of bacteria by macrophages; ii) the killing of phagocytized bacteria. In order to determine these processes, murine macrophage cells J774A.1 were exposed to bacteria (*Asaia*^WSP^ or *Asaia*^pHM4^) at different conditions (see Materials and Methods) of co-incubation at a MOI of 100 (100 bacteria:1 macrophage). After 1h of treatment with streptomycin, the intramacrophagic bacteria were quantified and expressed as CFU/ml. *Ad hoc* experiments showed that the uptakes, after 1h or 2h of incubation, were comparable (Supplementary Table 1a; p=0.823); for this reason the second time point (2h) was used in all the successive experiments, also considering published protocols (Migliore et al., 2018). Survival of bacteria after 24h of treatment was quantified and expressed as mean number of phagocytized bacteria in 2h and survived until 24h. As reported in Fig. 2a, the number of bacteria phagocytized by macrophages co-incubated with *Asaia*^WSP^ is higher than that of macrophages co-incubated with *Asaia*^pHM4^, and comparable with the uptake of *Asaia*^pHM4^ by macrophages treated with LPS (positive control): the mean number of bacteria *Asaia*^WSP^ phagocytized is almost the double than that of *Asaia*^pHM4^ (7.89×10^5^ CFU/ml vs 4.12×10^5^ CFU/ml). Bootstrap estimation confirmed that the increase in the number of phagocytized *Asaia*^WSP^ was statistically significant (Δ= 3.77×10^5^ CFU/ml, p=0.0006). As for the survival of bacteria in the macrophages, after 24h of co-infection most bacteria phagocytized during 2h of co-incubation are killed by the macrophages with slight differences between the two treated groups and the control. In particular, as reported in Supplementary Table 1b, a slightly lower number of bacteria *Asaia*^WSP^ were counted within macrophages (1.43×10^4^ CFU/ml), compared both to *Asaia*^pHM4^ and *Asaia*^pHM4^ + LPS (2.03×10^4^ CFU/ml and 1.07×10^4^ CFU/ml, respectively), but the differences are not statistically significant (p=0.628). The phagocytosis activity was also evaluated in pre-stimulated macrophages with *Asaia*, subsequently exposed to *S. epidermidis* for 1h or 2h. After both time points, the average number of phagocytized *S. epidermidis* after the pre-treatment with *Asaia*^WSP^ are higher than that of the controls, but with a slight overlap with the 0 intercepts (Supplementary Fig. 1).

**Fig. 2.**
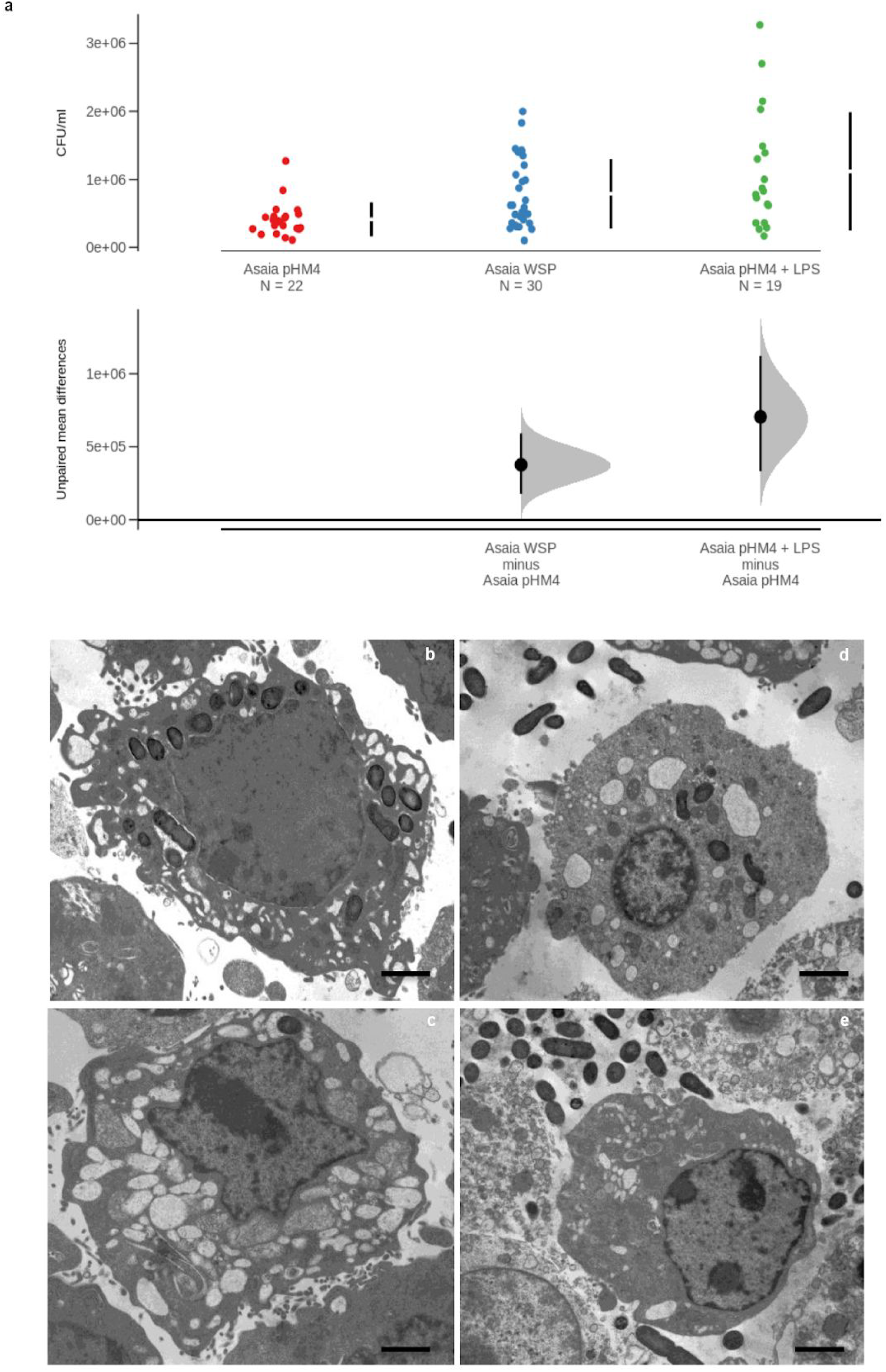
Bacterial uptake by macrophages and TEM observation. Murine macrophages were stimulated with *Asaia*^pHM4^, *Asaia*^WSP^, or *Asaia*^pHM4^ plus LPS. a) In the top panel of the estimation plot the numbers of bacteria internalized within macrophages are reported as CFU/ml. The vertical error bars denote mean and standard deviation of the observed data. In the bottom panel the unpaired mean differences between treatment groups compared to *Asaia*^pHM4^ are reported. The shaded distribution derives from the application of the resampling algorithm, the large black circle represents the average difference between groups, the error bars on the large black circle indicate the 95% confidence interval for the calculated difference. Estimation Statistics (ES) approach and bootstrapped Welch two-sample t-test were applied. The results are representative of triplicate experiments. b-e) TEM images of phagocytosis. Murine macrophages incubated with *Asaia*^WSP^ (b and c) or with *Asaia*^pHM4^ (d and e) for 24h. In panels b and c, note the bacteria in the vacuoles and the signs of intense phagocytic activity. In panels d and e, bacteria are mostly outside the macrophage. Bars: 5μm.

The intracellular localization of *Asaia*^WSP^ and *Asaia*^pHM4^ after 24h of incubation with macrophages was also investigated by transmission electron microscopy (TEM). As shown in Figs. 2b and c, in macrophages exposed to *Asaia*^WSP^, bacteria were observed inside the cells. In some cases, the phagocytic vacuole hosting the bacterium was evident; several empty vacuoles were also observed, which suggest that a strong digestive activity occurred in the treated macrophages. On the other hand, when the macrophages were treated with *Asaia*^pHM4^, the bacteria were mostly outside the macrophages and only a few bacteria were observed within the cells (Figs. 2d and e); in general, a reduced number of empty vacuoles were observed in the macrophages exposed to the control *Asaia*^pHM4^, compared to those exposed to *Asaia*^WSP^.

### Killing of *Leishmania* by *Asaia*^WSP^-stimulated macrophages

The above results encouraged us to investigate whether macrophage activation determined by *Asaia*^WSP^ could result in the phagocytosis and killing of *Leishmania*. The anti-leishmanial effect of *Asaia*^WSP^ was determined by microscopy observation after 24h and 48h of incubation, as reported in Maksouri et al. (2017). J774A.1 cells were pretreated with *Asaia* for 2h and then incubated with *L. infantum* parasites. The percentage of infected macrophages (infection rate) and the average numbers of *Leishmania* amastigotes in infected macrophages are reported, for each treatment group, in Supplementary Table 2a and b. The most informative time to evaluate an *in vitro* killing effect was 48h after *Leishmania* infection. Interestingly, based on the binomial negative count model (Fig. 3a and Supplementary Table 2c), the mean number of amastigotes in each macrophage significantly decreased from 1.61 (control group, macrophages exposed only to *Leishmania*) to 0.67 (*Asaia*^WSP^ + *Leishmania*), i.e. we observed a reduction of 58.4% (p<0.001). *Asaia*^pHM4^ also determined a slight reduction in amastigote number per macrophage (1.44) compared to *Leishmania* alone, but the difference was not significant (p=0.265). Moreover, a significant decrease of the mean number of *Leishmania* was recorded after the treatment with amphotericin B (0.45, p<0.001) (Supplementary Table 2c).

**Fig. 3.**
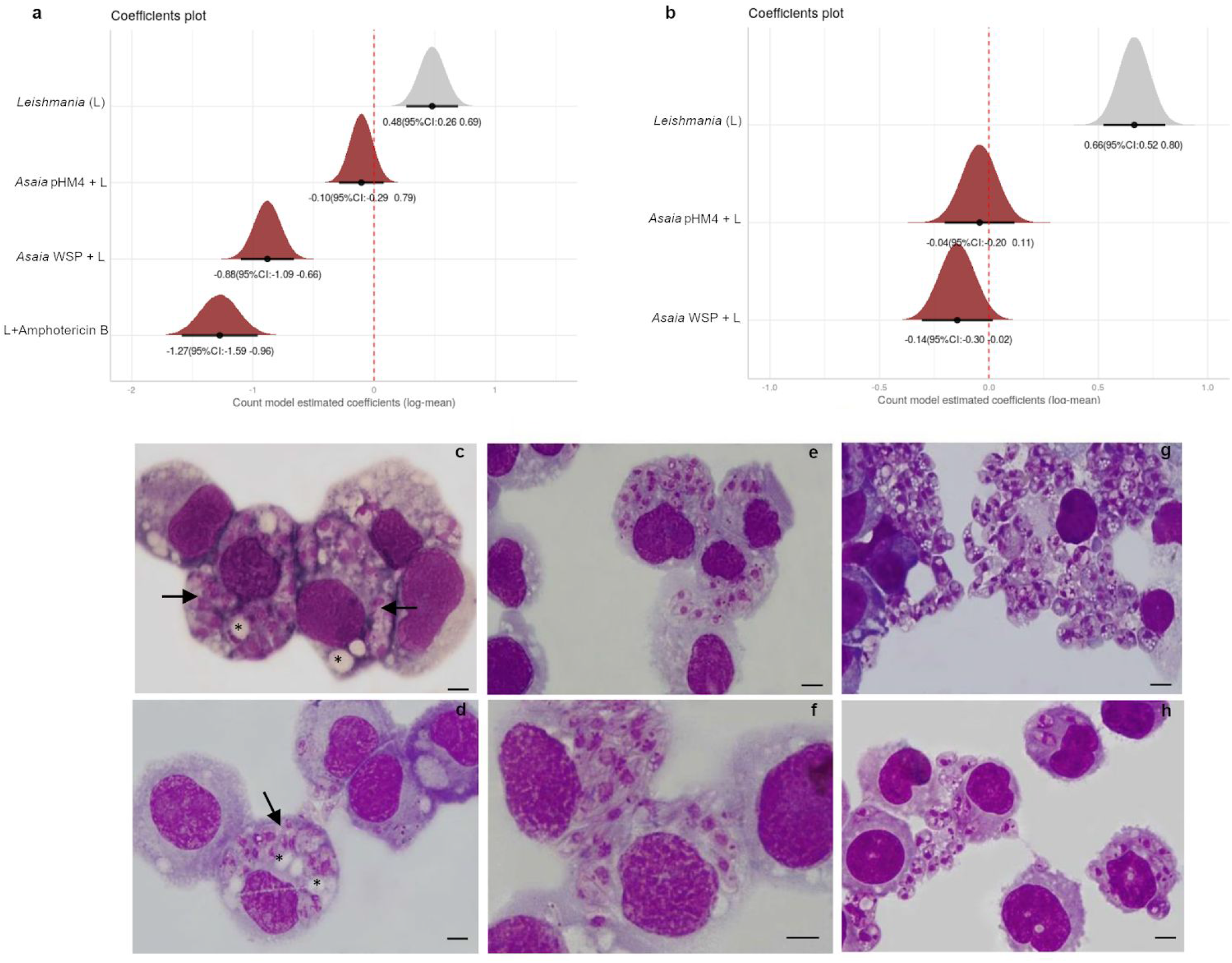
*Killing of* Leishmania *by* Asaia^WSP^*-stimulated macrophages after 48h of incubation.* Macrophages were pre-stimulated with *Asaia*^WSP^ or *Asaia*^pHM4^ or untreated, and then infected with *Leishmania* promastigotes (*Asaia*^WSP^ + L, *Asaia*^pHM4^ + L, *Leishmania* (L), respectively). Amphotericin B was used as control for the killing, for the observations at the 48h. a and b) Coefficients plots related to the 48h (a) or 24h (b) after *Leishmania* infections. Horizontal bars indicate the 95% confidence interval; vertical red lines correspond to the difference between the control group and the other groups. In panel a, the exponential of the estimated coefficient in group L (0.48) corresponds to the average number of amastigotes in this group. The other estimated coefficients (−0.10, −0.88, −1.27) correspond to the difference between the estimate of the reference group (L) and that of the treatment group. Similarly, in panel b, the exponential of the estimated coefficient in group L (0.66) corresponds to the average number of amastigotes in this group. The other estimated coefficients (−0.04, −0.14) correspond to the difference between the estimate of the reference group (L) and that of the treatment group. c-h) Giemsa staining of macrophages infected with *Leishmania* and *Asaia*^WSP^ (c and d), *Leishmania* and *Asaia*^pHM4^ (e and f) and *Leishmania* alone (g and h). Arrows indicate groups of killed amastigotes and asterisks indicate the presence of vacuoles in macrophages treated with *Asaia*^WSP^, signs of a high leishmanicidal activity. Bars: 5 μm. The experiments were performed in triplicate.

The Zero-inflated analysis in the ZINB model (Supplementary Table 2c) shows that the factor “type of treatment” was not significant as a predictor of the zero *Leishmania* counts in the macrophages (no significant differences were detected in the number of non infected macrophages in the different treatment groups compared to the control group). Figure 3 (panels c-h) shows the staining of macrophages co-infected with bacteria and *Leishmania*, after 48h. In panels c and d, infected macrophages present several vacuoles and the amastigotes show morphological changes e.g. loss of membrane integrity and formation of multiple cytoplasmic vacuoles. These cellular modifications are less noticeable in panels e and f which show the macrophages pre-treated with *Asaia*^pHM4^. Panels g and h (infection only with *Leishmania*) display numerous intact amastigotes, some of them in replication; part of the amastigotes is out of the macrophages.

Finally, the other time-point we analyzed was 24h. After 24h of infection, considering the analyses obtained applying the zero-inflated model and the count model, there were no significant differences between the groups. However, as for the results of the count model, we can observe that *Asaia*^WSP^ led to a reduction in the amastigote number per macrophage, with very limited overlap with the 0 intercept (Fig. 3b and Supplementary Table 2c).

### NO and ROS production by *Asaia*-stimulated macrophages

Two major effectors in macrophage microbial killing are NO and ROS (Fang, 2004), both involved in the killing of *Leishmania* parasites (Carneiro et al., 2016). These are produced by macrophages in resistant hosts, in response to *Leishmania* infection; which, on its side, tends to downregulate their production, especially in susceptible hosts (Ball et al., 2014). Considering the reduced survival of *Leishmania* within macrophages stimulated with *Asaia*^WSP^, we investigated whether this effect was associated with NO and ROS production. The capability of macrophages infected with *Asaia*^WSP^ or *Asaia*^WSP^ + *Leishmania* to produce NO was investigated, after 24h and 48h. After 24h post-infection, there are no significant differences between bacteria and controls, both in presence and in absence of *Leishmania* (Supplementary Fig. 2). Differently, Fig. 4a shows that, after 48h post-infection, the secretion of NO in the form of nitrites by macrophages treated with *Asaia*^WSP^ bacteria is statistically significant compared to the untreated macrophages (*Asaia*^WSP^= 132.71 μM, untreated macrophages = 77.11 μM; p=0.0038). *Asaia*^pHM4^ and *Leishmania* did not induce a significant increase in NO secretion compared to the unstimulated macrophages (*Asaia*^pHM4^= 80.46 μM, *Leishmania* = 78.10 μM; p=0.7092 and p=0.8687,respectively). Moreover, as shown in Figs. 4b and c, on detail, when the macrophages were incubated with *Asaia*^WSP^ the production of NO was significantly higher compared to *Asaia*^pHM4^, when the parasite is absent (*Asaia*^WSP^ w/o *Leishmania*= 132.71 μM; *Asaia*^pHM4^ w/o *Leishmania*= 80.46 μM; p=0.0002).

**Fig. 4.**
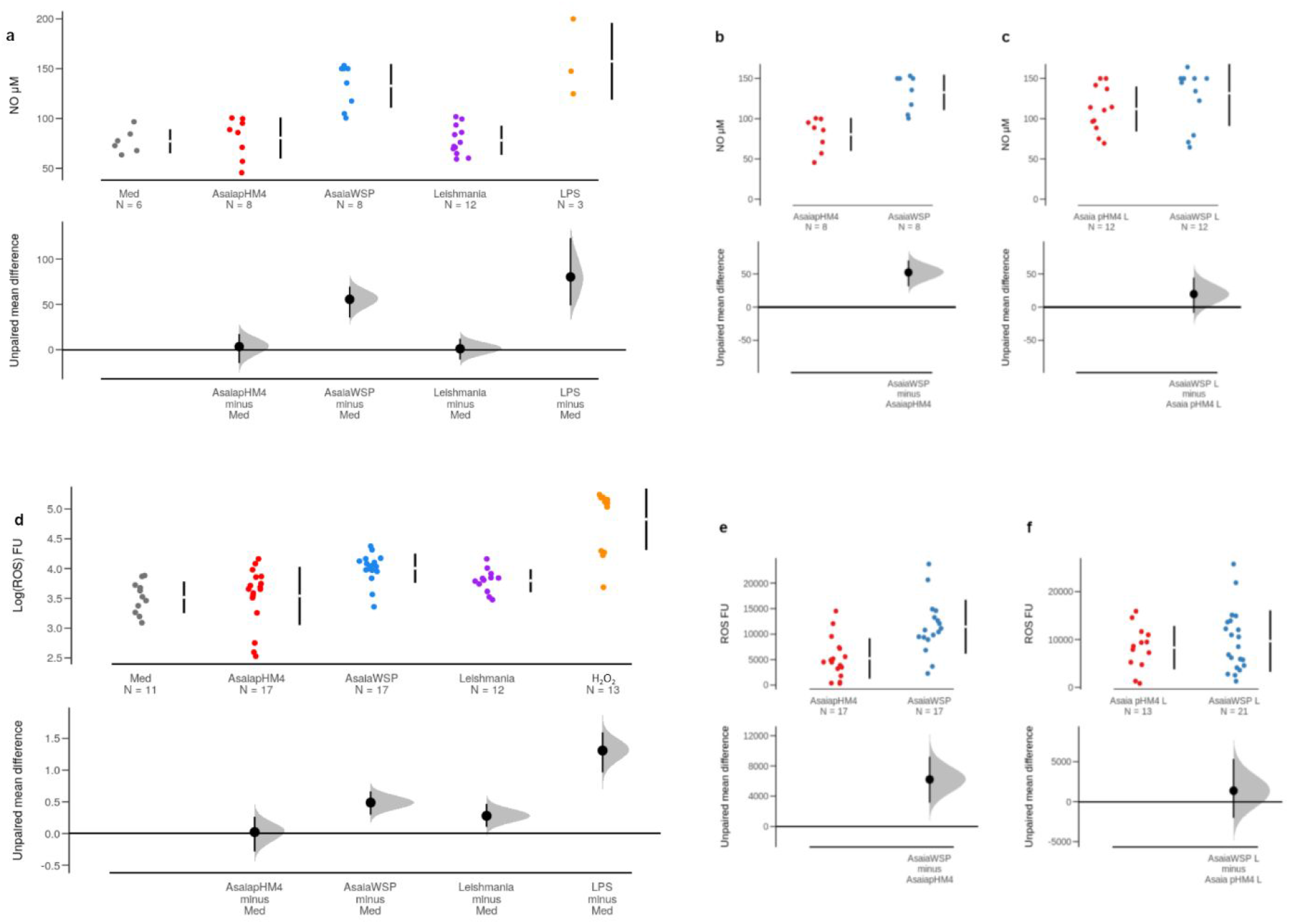
*Nitrites and ROS production by macrophages stimulated with* Asaia^WSP^. Nitrites and ROS levels were measured after 48h and 24h of incubation, respectively. a and d) The top panels of the estimation plot show the observed data of nitrites (μM) (a) and ROS (Log fluorescence unit, FU) (d) production by macrophages incubated with *Asaia*^pHM4^, *Asaia*^WSP^, *Leishmania*, positive control or untreated (Med). The vertical error bars denote mean and standard deviation of the observed data. In the bottom panels the unpaired mean differences between treatment groups compared to Med are reported. The shaded distribution derives from the application of the resampling algorithm, the large black circle represents the average difference between groups, the error bars on the large black circle indicate the 95% confidence interval for the calculated difference. b and c) Top panels show observed data of nitrites production by macrophages incubated with *Asaia*^WSP^ or *Asaia*^pHM4^ in absence of the parasite *Leishmania* (b) or in presence of the parasite (c). Bottom panels show the the comparisons between *Asaia*^WSP^ and the control bacterium *Asaia*^pHM4^. e and f) FU released by macrophages incubated with *Asaia*^WSP^ or *Asaia*^pHM4^ w/o *Leishmania* (e) or with *Leishmania* (f) at the top and the unpaired mean comparisons between the two bacteria at the bottom; Estimation Statistics (ES) approach and bootstrapped Welch two-sample t-test were applied. The results are representative of triplicate experiments.

NO status was also assessed by measuring inducible NO synthase (iNOS). The cells were collected and analysed for the expression of iNOS gene, at the first time point (24h) by reverse transcription-quantitative PCR. As reported in Supplementary Table 3a, iNOS relative expression after exposure to *Asaia*^WSP^ was higher than that of cells infected with *Leishmania* or *Asaia*^pHM4^, supporting what we observed analysing NO production (*Asaia*^WSP^= 1.35, *Asaia*^pHM4^ = 0.2, *Leishmania* = 0.002, untreated = 0.05, LPS = 3.04). In addition to nitrites, macrophages in response to microbial infection produce reactive oxygen species (ROS) as a killing mechanism, in the context of the M1 phenotype. The production of ROS after 24h in macrophages was investigated by a fluorometric assay. In Fig. 4d bootstrap estimation analyses show the significant production of ROS determined by the treatment of macrophages with *Asaia*^WSP^ (4 Log FU) or with LPS (4.82 Log FU), compared with untreated macrophages, as control (3.51 Log FU) (p=0.0002 and p=0.0002, respectively). Interestingly, *Leishmania* alone (3.79 Log FU) determined an appreciable increase of ROS compared to untreated macrophages (p=0.008). As reported in Figs. 4e and f, when macrophages were stimulated by *Asaia*^WSP^ the production of ROS is significantly different compared to the *Asaia*^pHM4^, when the parasite is absent (*Asaia*^WSP^ w/o *Leishmania*= 11454 FU; *Asaia*^pHM4^ w/o *Leishmania*= 5237 FU; p=0.0006).

### M1 cytokines secretion by *Asaia*-stimulated macrophages

Macrophages were stimulated with *Asaia* (MOI 100) or with *Asaia* bacteria plus *Leishmania* (2 *Leishmania*:1 macrophage, MOI 2) promastigotes; after 24h and 48h of co-infection, the culture supernatants were collected and analyzed by ELISA assay, for the secretion of the cytokines TNFα, IL-12p40, IL-1β and IL-6, as markers of M1 polarization (Fig. 5; for a complete description of the choice of cytokines see Supplementary Text). Twenty-four hours post-infection, the production of TNFα cytokine by macrophages treated with both *Asaia* bacteria was statistically different compared to the untreated macrophages, indicating a strong effect due to the presence of the bacteria (*Asaia*^WSP^= 2.95 Log pg/ml, p<0.001; *Asaia*^pHM4^= 2.88 Log pg/ml, p<0.001; untreated macrophages= 0.76 Log pg/ml; Fig. 5a). Comparing the effect of the two bacteria, we can observe (Figs. 5b and c) a higher production of this cytokine by *Asaia*^WSP^ compared to the *Asaia*^pHM4^ treated cells, both in the presence or in the absence of the *Leishmania,* although there was no significant difference between the groups. The same trend of differences was observed for the secretion TNFα cytokine after 48h post-infection (Supplementary Fig. 3). The same tendency was obtained for the expression of the cytokine IL-12p40, 24h post-infection (Figs. 5d-f). The macrophages treated by *Asaia*^WSP^ and *Asaia*^pHM4^ produced a greater amount of cytokine IL-12p40 comparable with the production determined by LPS treatment (*Asaia*^WSP^= 3.74 Log pg/ml, *Asaia*^pHM4^= 3.63 Log pg/ml, LPS = 3.46 Log pg/ml) and statistically higher than the control (0.72 Log pg/ml) (*Asaia*^WSP^: p<0.001; *Asaia*^pHM4^: p<0.001; LPS: p<0.001). *Leishmania* did not statistically affect the production of this cytokine (0.50 Log pg/ml, Fig. 5d). As observed for TNFα, we obtained a higher production of IL-12p40 by *Asaia*^WSP^ compared to the *Asaia*^pHM4^ treated cells (i.e. with differences between the sample means), though there was no significant difference between the groups (Figs. 5e and f). Culture supernatants from stimulated macrophages were also checked for the production of the cytokines IL-1β (Figs. 5g-i) and IL-6 (Figs. 5j-l). The secretion of IL-1β cytokine by macrophages treated with both *Asaia* bacteria was statistically significant compared to untreated macrophages (*Asaia*^WSP^: 2.55 Log pg/ml, p<0.001; *Asaia*^pHM4^: 2.47 Log pg/ml, p<0.001). Noteworthy, the unpaired mean differences of macrophages treated with both bacteria are higher than that of macrophages treated by LPS (2.08 Log pg/ml) (Fig. 5g). However, the unpaired mean comparison of IL-1β of *Asaia*^WSP^ vs *Asaia*^phM4^ was not significant (Figs. 5h and i). Finally, the quantification of IL-6 induced by *Asaia*^WSP^ and *Asaia*^pHM4^ after 24h was comparable with those of LPS treated macrophages and higher than that of the control (*Asaia*^WSP^= 3.74 Log pg/ml, *Asaia*^pHM4^= 3.60 Log pg/ml, LPS = 4.35 Log pg/ml; untreated macrophages = 0.99 Log pg/ml; p<0.001 for all comparisons. Fig. 5j). As reported in Figs. 5k and l, when the macrophages were infected by *Asaia*^WSP^ the production of IL-6 is significantly higher compared to the *Asaia*^pHM4^; bootstrap analysis confirmed the significance when the parasite was absent (p=0.0012).

**Fig. 5.**
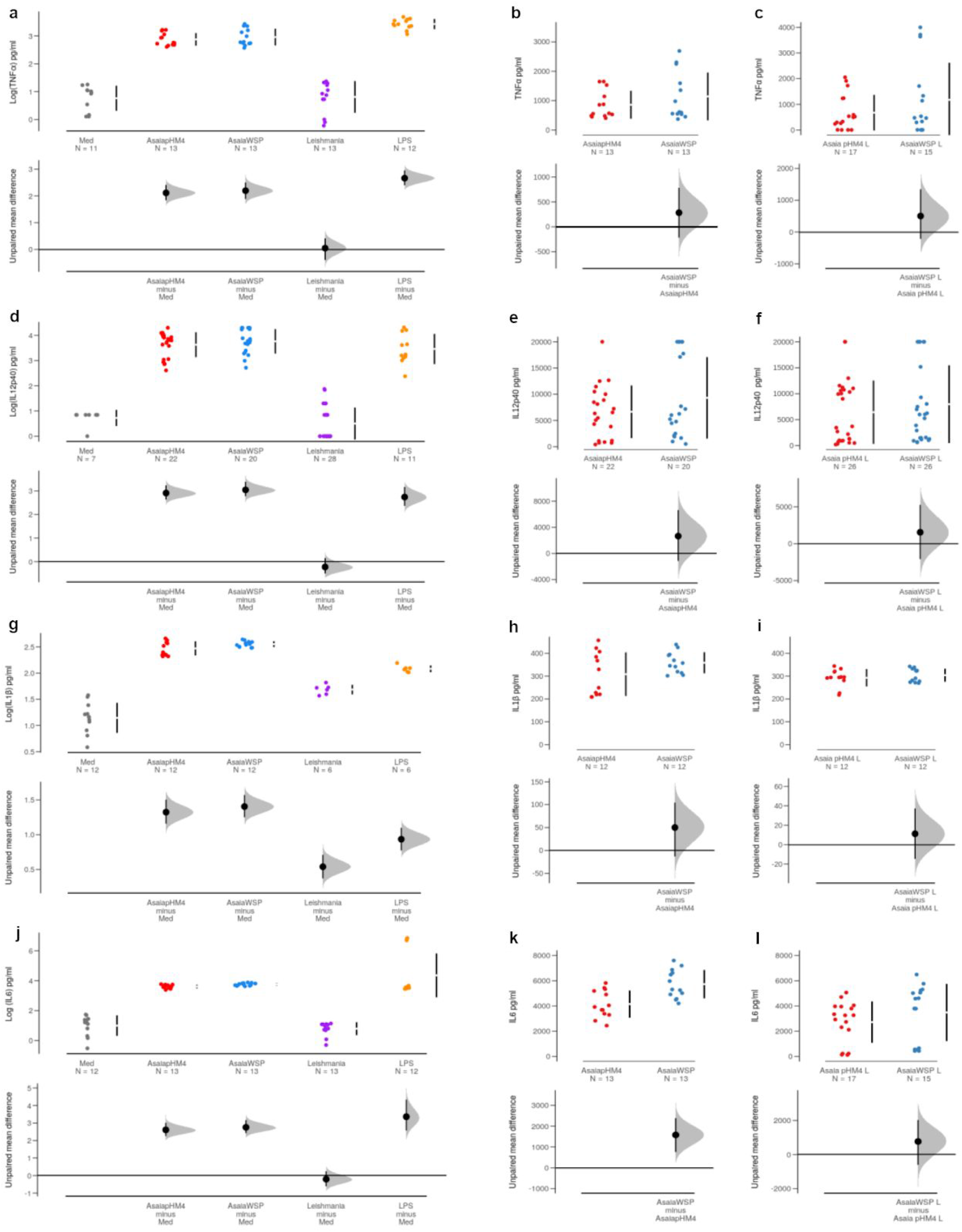
*TNFα, IL-12p40, IL-1β and IL-6 cytokines produced by macrophages stimulated with* Asaia *bacteria.* Levels of TNFα (a), IL-12p40 (d), IL-1β (g) and IL-6 (j) cytokines expressed as Log pg/ml and produced by macrophages treated with *Asaia*^WSP^, *Asaia*^pHM4^, *Leishmania*, LPS or untreated (Med) after 24h of co-incubation are represented in the top panels. The vertical error bars denote mean and standard deviation of the observed data. In bottom panels the unpaired mean comparisons of TNFα (a), IL-12p40 (d), IL-1β (g) and IL-6 (j) between treatment groups and untreated macrophages, as control group, are shown. The shaded distribution derives from the application of the resampling algorithm, the large black circle represents the average difference between groups, the error bars on the large black circle indicate the 95% confidence interval for the calculated difference. b and c) Top panels report observed data of TNFα production by macrophages incubated with *Asaia*^WSP^ or *Asaia*^pHM4^ in absence of the parasite *Leishmania* (b) or in presence of the parasite (c). Bottom panels show the comparisons between *Asaia*^WSP^ and the control bacterium *Asaia*^pHM4^. e and f) IL-12p40 produced by macrophages incubated with *Asaia*^WSP^ or *Asaia*^pHM4^ w/o *Leishmania* (e) or with *Leishmania* (f) at the top and the unpaired mean differences between the two bacteria at the bottom. h and i) Top panels report observed data of IL-1β production by macrophages incubated with *Asaia*^WSP^ or *Asaia*^pHM4^ in absence of the parasite *Leishmania* (h) or in presence of the parasite (i). Bottom panels show the comparisons between *Asaia*^WSP^ and the control bacterium *Asaia*^pHM4^. k and l) IL-6 produced by macrophages incubated with *Asaia*^WSP^ or *Asaia*^pHM4^ w/o *Leishmania* (k) or with *Leishmania* (l) at the top and the unpaired mean differences between the two bacteria at the bottom. Estimation Statistics (ES) approach and bootstrapped Welch two-sample t-test were applied. Observed data are representative of three independent experiments.

### M2 cytokine secretion and expression of *arginase I*

Culture supernatants from cells pre-stimulated with LPS and infected with *Asaia* bacteria or *Asaia* bacteria + *Leishmania* were tested for the production of IL-10, a typical marker of M2 polarization. In Supplementary Table 3b we reported the means, the standard deviation and the corresponding p-value. We observed that macrophages incubated with *Asaia*^pHM4^ produced more IL-10 compared to the *Asaia*^WSP^ when the cells were infected only with bacteria (*Asaia*^WSP^ w/o *Leishmania*= 565.81 pg/ml; *Asaia*^pHM4^ w/o *Leishmania*= 692.05 pg/ml). As for *arginase I* expression, cells were analyzed at 48h. As shown in Supplementary Table 3c when macrophages were exposed to *Asaia*^WSP^ in presence of *Leishmania,* there was a downregulation of the gene expression, compared to the *Asaia*^pHM4^ treatment (*Asaia*^WSP^ + *Leishmania*= 0.37; *Asaia*^pHM4^ + *Leishmania*= 0.76).

### Costimulatory molecules and MHC-II expression by *Asaia*-stimulated macrophages

To investigate the effect of *Asaia* on the expression of selected surface markers (CD80-CD86-CD40) involved during *Leishmania* infection, after 24h of infection with bacteria and *Leishmania,* macrophages were processed for flow cytometry analyses. The percentage of cells pretreated with *Asaia*^WSP^ or *Asaia*^pHM4^ and expressing CD40 marker, was comparable (i.e. no statistical differences were observed) to that of macrophages treated with LPS, the positive control (LPS: 79%; *Asaia*^WSP^: 70%; *Asaia*^pHM4^: 58%) (Fig. 6). The expression of CD40 on untreated macrophages or on macrophages infected only with *Leishmania* was statistically lower than the LPS positive control (untreated macrophages: 8%, *Leishmania*: 18%, p<0.001 for both comparisons). The same trend was obtained from CD80 and CD86 analyses. For both markers, pre-treatments with *Asaia*^WSP^ or *Asaia*^pHM4^ stimulated a percentage of macrophages to present these molecules comparable with the positive control (CD80^+^ cells: LPS= 33%, *Asaia*^WSP^= 22%, *Asaia*^pHM4^= 17%; CD86^+^ cells: LPS= 59%, *Asaia*^WSP^= 40%, *Asaia*^pHM4^= 34%). On the contrary, the proportion of positive cells in untreated macrophages or in macrophages infected only with *Leishmania* was statistically lower than LPS (CD80^+^ cells: untreated macrophages= 9% p=0.008, *Leishmania*: 8%, p=0.003; CD86^+^ cells: untreated macrophages= 4% p<0.001, *Leishmania*= 15% p=0.001; Supplementary Table 4 and Fig. 6). Fig. 6 also shows the expression of MHC-II by macrophages infected with bacteria and *Leishmania* after 48h of infection and pre-stimulated with IFNγ. The priming with IFNγ was necessary to stimulate the expression of MHC-II by this cell line, because the constitutive expression on J774A.1 is very low (Kalupahana et al., 2005). The proportion of MHC-II positive cells treated with *Asaia*^WSP^ was statistically higher than the positive control treatment with PMA (Phorbol 12-Myristate 13-Acetate) (*Asaia*^WSP^: 43%, PMA: 30%, p= 0.027; Supplementary Table 4). The group *Asaia*^pHM4^ showed a higher proportion of positive cells compared with the positive control, but not statistically significant (p= 0.545). On the contrary, the proportion of positive cells in the untreated group was low (4.3%) and statistically significant compared with the positive control (p<0.001; Supplementary Table 4).

**Fig. 6.**
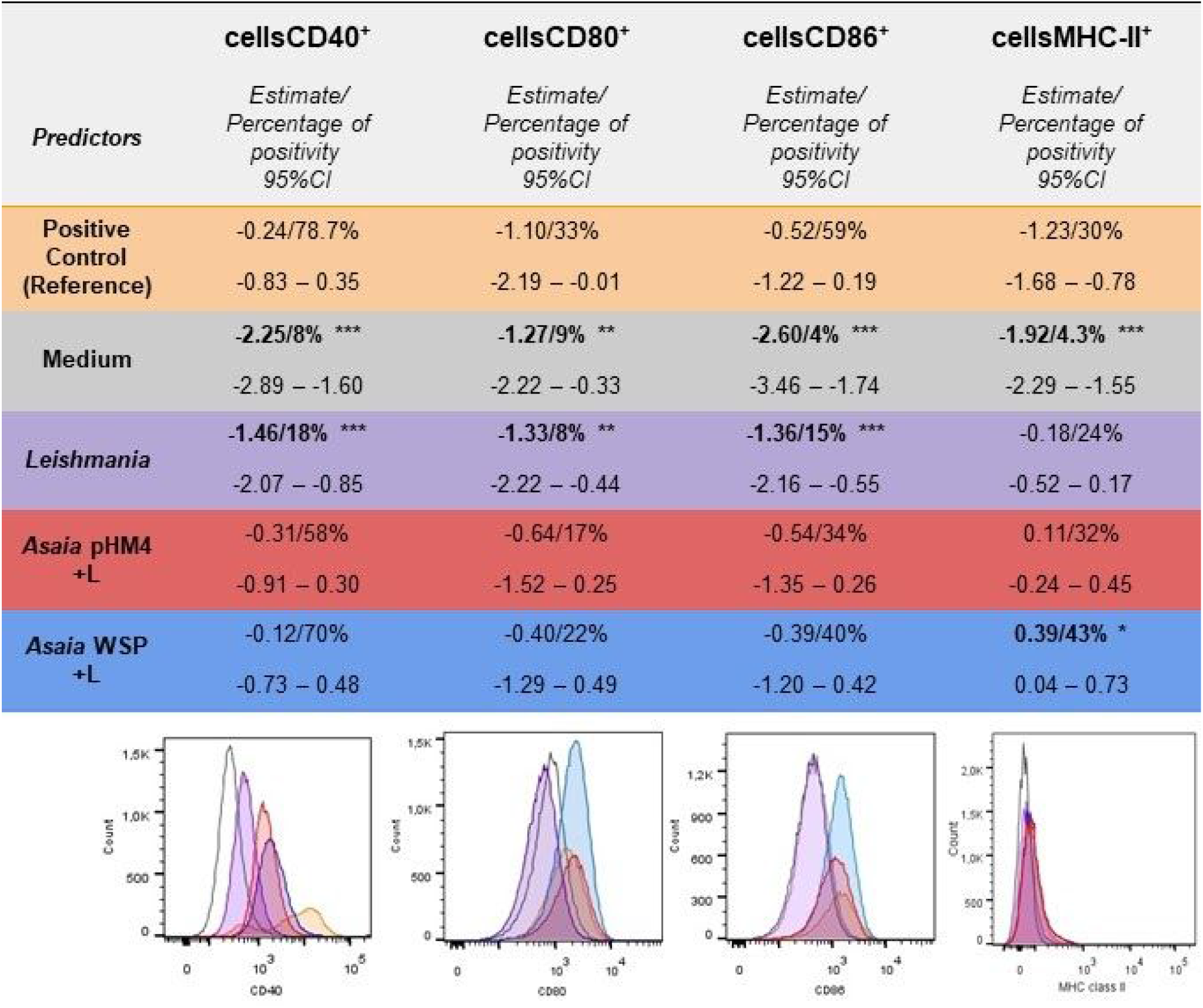
*Costimulatory molecules and MHC-II expression by macrophages stimulated with* Asaia *bacteria.* a) Differences in the proportions of macrophages positive for CD40, CD80 and CD86 costimulatory molecules and MHC-II marker between different treatment groups are reported. e^(intercept+estimated coefficient)^ can be interpreted as the proportion of cells in the different treatment groups. b) Flow cytometry histograms of CD40, CD80 and CD86 and MHC-II expression, representative of three independent experiments are showed (orange curve: positive control; grey curve: negative control; violet curve: *Leishmania*; red curve: *Asaia*^pHM4^ + L; blue curve: *Asaia*^WSP^ + L).

## Discussion

Our main goal was to determine whether *Asaia*^WSP^ is capable to induce a macrophage activation, strong enough to result in microbicidal effects, particularly on *Leishmania*. A first set of assays showed that *Asaia*^WSP^ indeed activates an antibacterial response in macrophages, on both *Asaia* itself and on *S. epidermidis*. The study was then focused on *Leishmania*, an intracellular parasite that targets monocytes and macrophages (Antoniou et al., 2013; Rossi & Fasel, 2018). The experiments were realized using a macrophage cell line derived from a *Leishmania*-susceptible mice strain. In susceptible hosts, macrophages are typically anergic towards *Leishmania*, which replicates in these cells (Nigg et al., 2007). As expected, our experiment showed that macrophages infected with *Leishmania* alone, with no pre-stimuli, reached a high load of parasites. However, after *Asaia*^WSP^ pre-stimulation, *Leishmania* survival was dramatically reduced, with an effect that approximated that of amphotericin B, a choice drug to achieve the killing of *Leishmania* parasites. On the other hand, a higher amastigote survival was observed after pre-stimulation with the control bacterium *Asaia*^pHM4^, which indicates that the stronger effect determined by *Asaia*^WSP^ is likely associated with the expression of WSP. In summary, *Asaia* expressing WSP is capable to revert the anergy of macrophages from a *Leishmania* susceptible host, restoring their capability to mount an effective microbicidal response. Indeed, the strong reduction in the number of *Leishmania* amastigotes determined by *Asaia*^WSP^ likely derives from the activation of the killing activity of macrophage cells. At 48h after *Asaia*^WSP^ stimulation the number of amastigotes per macrophage was significantly lower compared to the number observed at 24h. In particular, comparing the results obtained at 24h and 48h after *Asaia*^WSP^ stimulation, the average number of amastigotes per macrophage decreased passing from 1.68 to 0.67 (60% of reduction), contrary to macrophages infected with *Leishmania* alone, in which the number of amastigotes changed only slightly (from 1.93 to 1.61, 17% of reduction). Microscopic observations are coherent with the occurrence of leishmanicidal activity in *Asaia*^WSP^-stimulated macrophages: amastigotes showed degeneration of the membranes and macrophages displayed several vacuoles, signs of an intense digestive activity. Indeed, in our experiments *Asaia*^WSP^ determined a significant increase in the production NO and ROS, two major effectors in macrophage anti-bacterial and anti-*Leishmania* responses (Fang, 2004; Carneiro et al., 2016). The increased killing activity after *Asaia*^WSP^ stimulation can thus be attributed to a classical M1 macrophage activation, with production of well-established microbicidal effectors. NO production is governed by the inducible NO synthase (iNOS), whose expression was actually upregulated after macrophage stimulation with *Asaia*^WSP^. iNOS, a typical marker of M1 macrophage activation, shares the same substrate with the enzyme arginase, whose upregulation is instead crucial in the process of the alternative, M2 macrophage polarization (Biswas et al., 2012). The arginase assay revealed that *Asaia*^WSP^ reduces the expression of this M2-associated enzyme, in the presence of *Leishmania*. Furthermore, assays on cytokine production revealed that both *Asaia*^WSP^ and *Asaia*^pHM4^ induced overexpression of most of the tested M1 cytokines, with a slight, but coherent, overexpression of these cytokines after stimulation with *Asaia*^WSP^ (for a discussion about the choice of the cytokines to be quantified and their role in leishmaniases, see Supplementary Text). As for type-2 cytokines, IL-10 was significantly downregulated by *Asaia*^WSP^ in the presence of *Leishmania*. In summary, the activation of macrophage phagocytosis and killing activity, the induction of NO and ROS production, and the opposed effects on the expression of M1 and M2 markers (iNOS, arginase and cytokines) coherently point to the capability of *Asaia*^WSP^ to activate macrophages, with a bias towards the classical, M1 phenotype. *Asaia*^WSP^ proved to be very effective also in the induction IL-6, a pro-inflammatory cytokine that however does not perfectly fits into the Th1/Th2 paradigm (Diehl & Rincón, 2002).

During the immune response, activation of T CD4^+^ lymphocytes by macrophages requires antigen presentation through MHC-II, and the concomitant expression of co-stimulatory molecules, which also play a role in immune-modulation (Podojil & Miller, 2009; for a discussion about the choice of co-stimulatory molecules and their role in leishmaniases, see Supplementary Text). The analysis of the expression of co-stimulatory molecules showed that pre-treatment with *Asaia*^WSP^ or *Asaia*^pHM4^ stimulated a higher number of macrophages to present the tested CD receptors, compared to the control (Fig. 6). *Asaia* bacteria also stimulated the expression of MHC-II, an effect that was higher in cells pretreated with *Asaia*^WSP^, compared to the PMA control treatment (Fig. 6). The evidence that *Asaia* stimulates the expression of both MHC-II and co-stimulatory molecules highlights the potential of these bacteria to determine a complete macrophage activation, with expression of the molecules required for antigen presentation and for the co-stimulation of T CD4^+^ cells.

Our study was not aimed at the investigation of the immune response to *Leishmania* parasites, a deeply explored field (e.g. Gupta et al., 2013). However, we emphasize that the results of our experiments on *Leishmania* alone confirmed the ability of this parasite to determine low level of M1 activation, limited expression of MHC-II and co-stimulatory molecules, and increased expression of M2 markers, in agreement with its well-documented adaptation to intra-macrophage survival and multiplication (e.g. Zutshi et al., 2019).

In this study, the stronger capability of *Asaia*^WSP^ to induce an effective macrophage activation in comparison with *Asaia*^pHM4^, as revealed by the *Leishmania* killing assay, provides sound evidence for the capacity of WSP to determine macrophage activation, with clear signs for M1 polarization. Therefore, WSP from the *Wolbachia* endosymbionts of filarial nematodes is confirmed as a candidate immune-modulating factor in filarial diseases, in agreement with the suggestions of Brattig et al. (2004). Interestingly, preinfection of hamsters with *Brugia malayi* (a *Wolbachia*-harbouring filaria) determined significant protection against a successive challenge with *Leishmania donovani* (Murthy et al., 2008). In addition, epidemiological studies have so far revealed a limited number of cases of co-infection in dogs, with *Dirofilaria* and *Leishmania* (Medkour et al., 2020; Gizzarelli et al., 2019). On the other hand, in dogs co-infected by *D. immitis* and *L. infantum* the presence of circulating *Wolbachia* DNA negatively correlates with clinical signs of leishmaniasis (Maia et al., 2016). We could thus speculate that in dogs infected by the blood-dwelling filaria *D. immitis* the continuous release of WSP-loaded *Wolbachia* into the blood stream could modulate the immunity of the host, protecting it from *Leishmania* infection (even though chronic filarial infections in humans is also associated with macrophage tolerization; e.g. see Hoerauf et al., 2005).

A first issue addressed by this study was whether WSP, the *Wolbachia* surface protein, possesses the capability to activate macrophages, determining an increased microbicidal activity. The second main issue was whether the engineered bacterium *Asaia*^WSP^ could be used to induce macrophage activation and *Leishmania* killing. Our results, while providing a positive answer to both the above questions, revealed that also the tested control bacterium, *Asaia*^pHM4^, displays macrophage activating properties. The fact that both *Asaia* strains are able to activate macrophages is not surprising, considering their bacterial Gram-negative nature. It is interesting that *Acetobacter pasteurianus*, a member of the *Acetobacteraceae*, phylogenetically related with the genus *Asaia*, has recently been shown to induce macrophage activation in the same cell line used in this study, through the release of micro-vesicles (Hashimoto, 2018).

With few exceptions, *Leishmania* parasites are paradigmatic for their capability to skew macrophages into an anergic, M2 state (Tomiotto-Pellissier et al., 2018). Our study reveals that both tested *Asaia* bacteria, but particularly *Asaia*^WSP^, possess the ability to reverse this anergic state, towards a classical M1 activation. This suggest that *Asaia*, either naïve or engineered, is worth to be investigated as a potential prophylactic or therapeutic agent, for the control of leishmaniases and other M1/Th1-impaired diseases. The M1-polarizing properties of *Asaia* could be exploited without further manipulations, or after its engineering for the expression of specific antigens, from *Leishmania* or other pathogens. Based on the results here presented, *Asaia*^WSP^ appears as the most promising, considering its ability to induce the killing of *Leishmania*. The effective internalization of *Asaia*^WSP^ in macrophages, followed by proper M1 activation, including expression of MHC-II and costimulatory molecules, suggest that this bacterium, once further modified for antigen expression, holds the potential to stimulate specific T CD4^+^ lymphocytes as Th1-polarized memory cells. In prospects, the eventual use of *Asaia* bacteria as engineered vehicles for immunomodulation or vaccination will require that safety issues are properly addressed. As a general conclusion we highlight that *Asaia* bacteria prove to be suitable for genetic manipulation, safe and easy to handle (Epis et al., 2012), and capable of inducing activation of macrophages, with a killing *Leishmania* parasites that is significantly potentiated by the expression of the *Wolbachia* surface protein.

## Materials and methods

### Cell and parasite cultures

The J774A.1 ATCC® TIB-67 macrophage cell line, derived from *Leishmania*-susceptible BALB/c mice, was grown in Dulbecco’s Modified Eagle’s Medium (DMEM) supplemented with 10% fetal bovine serum (FBS) and maintained under an atmosphere of 5% CO_2_ at 37°C in incubator. All reagents for the cell cultures were purchased from ATCC (Manassas, VA, USA). The *Leishmania infantum* promastigotes derived from a strain maintained at the Istituto Superiore di Sanità, Rome, Italy (strain MHOM/TN/80/IPT1). The parasites were grown at 23°C in Schneider’s *Drosophila* medium (Thermo Fisher Scientific, Waltham, MA) supplemented with 10% FBS and gentamycin (5 μg/ml).

### Bacterial strains and growth conditions

The bacterium *Asaia*^WSP^ derived from the engineering of the bacterium *Asaia* SF2.1 strain, originally isolated from the *Anopheles stephensi* mosquito (Favia et al., 2007), with the plasmid pHM4-WSP (Epis et al., 2020). Briefly, the WSP cassette, inserted in the plasmid pHM4, is composed by the *Wolbachia* surface protein gene sequence (from the *Wolbachia* of the nematode *Dirofilaria immitis*), the neomycin phosphotranferase promoter PnptII, the E-TAG epitope and the transcription terminator Trrn. *Asaia*^pHM4^ was also obtained from strain *Asaia* SF2.1, but transformed with the empty plasmid (without the WSP-coding gene) and was used as control bacterium (Epis et al., 2020). Both bacteria were grown overnight in GLY medium broth (glycerol 25 g/l and yeast extract 10 g/l, pH 5) supplemented with kanamycin 100 μg/ml, under constant agitation at 30°C.

### Phagocytosis assays on bacteria and Transmission Electron Microscopy

Macrophages were seeded in 24-well plates (2×10^5^/ml) and allowed to adhere overnight at 37°C in humidified 5% CO_2_ atmosphere. Phagocytosis assay was performed applying the gentamicin protection assay as reported in Glasser et al., 2001, with minor modifications. *Asaia* bacteria were washed with sterile PBS and resuspended in complete DMEM medium. The macrophages were treated at a multiplicity of infection (MOI) of 100 bacteria per macrophage. Macrophages were co-incubated with *Asaia*^pHM4^ or *Asaia*^WSP^; as a positive control, macrophages were infected with *Asaia*^pHM4^ in presence of *Escherichia coli* lipopolysaccharide (LPS) (0.3 μg/ml) (R&D Systems, Minneapolis, MN). After a 10 min centrifugation at 1,000 rpm, macrophages were incubated 1h or 2h at 37°C to allow internalization of the bacteria (Migliore et al., 2018). Then, the macrophages were washed and treated with complete DMEM containing 100 μg/ml streptomycin for 1h at 37°C to kill extracellular bacteria. After two washes with PBS, part of the macrophages was lysed using deionized water containing 1% (vol/vol) Triton X-100 (Sigma Aldrich, USA) for 15 min at 37°C, to release phagocytized bacteria. The bacterial titre was determined by plating ten-fold serial dilutions of the cell lysates on GLY plates and CFU/ml were counted after growth for 48h at 30°C. In addition, to determine the bacterial survival inside the cells, the remaining part of the macrophages, after the treatment with streptomycin 100 μg/ml, were incubated with streptomycin 20 μg/ml until 24h of infection, followed by the final step of the protocol, as described above.

As for phagocytosis data analysis after the assays on *Asaia* bacteria, all data were stored in a spreadsheet and then analysed according to a one-way factor design with “Treatment” as the main experimental factor of interest. For experiments testing the effect of the “Treatment” (*Asaia*^WSP^, *Asaia*^pHM4^, *Leishmania*, LPS vs Med) the Estimation Statistics (ES) approach was applied (Wheaton, 2012). ES is a simple framework that, while avoiding the pitfalls of significance testing, uses familiar statistical concepts: means, mean differences, and error bars. More importantly, it focuses on the effect size of one’s experiment/intervention, as opposed to significance testing by calculating effect size (mean differences) with his 95% confidence interval, using the bias-corrected and accelerated (BCa) bootstrap confidence interval of Efron and Tibshirani (1993; Efron, 1981) and suggested by Rózsa et al., 2000. The ES approach produces a plot which presents the raw data (top panel) and the bootstrap confidence interval of the effect size (the difference in means, bottom panel) aligned as a single integrated plot (e.g. see Fig. 2). However, in order to quantify the strength of evidence against null hypothesis (mean difference = 0), Fisher’s significant test was applied (bootstrapped Welch two-sample t-test) and the exact p-values are reported. Data analysis were performed in R language using dabestr package for ES (bias-corrected and accelerated (BCa) bootstrap confidence interval and Cumming plot) and Mkinfer package for bootstrapped Welch two-sample t-test, boot.t.test function (see Supplementary Information for supplementary references).

Efficiency of the phagocytosis was evaluated also against the bacterium *Staphylococcus epidermidis* (ATCC 155) following the protocol described above. *S. epidermidis* was grown in LB medium broth buffered to 7.0-7.4 pH with NaOH under constant rotation at 37°C, overnight. Briefly, macrophages were first incubated with *Asaia* for 2h (100 bacteria:1 macrophage, MOI 100) and then were incubated with *S. epidermidis* (10 bacteria:1 macrophage, MOI 10) for 1h or 2h at 37°C. The protocol described above was then applied. The analyses of the data were performed applying the ES approach, as deeply described above. The *Asaia* uptake by macrophages was also evaluated by TEM. After 24h of infection, cells were pelleted, washed with PBS and immediately fixed in 0.1 M cacodylate buffer (pH 7.2) containing 2.5% glutaraldehyde for 2h at 4°C and postfixed in 1% O_s_O_4_ in 0.1 M cacodylate buffer (pH 7.2) for 1.5h at 4°C. Subsequently, the samples were subject to dehydration in ethanol and then were embedded in Epon 812. Finally, thin sections were stained with uranyl acetate and lead citrate and examined under an EM900 TEM (Zeiss).

### *Leishmania* infection assay and killing determination

Macrophages were seeded in 6-well plates (2×10^5^/ml) and allowed to adhere overnight at 37°C in humidified 5% CO_2_ atmosphere. The macrophages were stimulated with *Asaia*^pHM4^ or *Asaia*^WSP^ at a MOI of 100 bacteria per macrophage for 2h and then treated with streptomycin for 1h. Subsequently, cell monolayers were washed once in PBS and then infected with *L. infantum* stationary phase promastigotes at a ratio of 2:1 (2 parasites per 1 macrophage). Non-internalized promastigotes were removed at 24h post infection by washing with PBS and fresh DMEM was replaced. Cells were then maintained at 37°C for further 24h (for a total of 48h from the infection). At designated time points (24h and 48h) the culture supernatants were collected and stored until determination of cytokines and nitrites (see below). For the assessment of leishmanicidal activity, macrophages after 48h of treatment were washed with PBS, collected by using a cell scraper, centrifugated at 1,200 rpm for 6 min and washed with PBS. Finally, they were suspended in 200 μl PBS at the final concentration of 10^6^cells/ml and cytocentrifugated (Cytospin, Hettich) for 5 min at 500 rpm on a slide and stained with Giemsa solution following the standard protocol (Sigma-Aldrich, USA). As control of the leishmanicidal activity, we treated the macrophages with the anti-*Leishmania* drug amphotericin B (0.3 μg/ml). The number of parasites in infected macrophages and the percentage of infected macrophages (infection rate) were determined with a microscope at 100X; ten areas of two slides per treatment were used to determine these indices. The experiments were performed in triplicate.

The association between the treatment factor (*Asaia*^WSP^, *Asaia*^pHM4^, amphotericin B, *Leishmania* alone) and the count outcome, i.e. the number of amastigotes in the macrophages, was determined using a zero-inflated negative binomial (ZINB) model (Islam et al., 2019; see Supplementary Information for supplementary references), since the recorded data were overdispersed and with an excess of zeros (we indeed recorded an overall 63% macrophages with zero amastigotes in the four groups). The amastigote number in the macrophages infected with *Leishmania* alone was used as the reference. A ZINB model assumes that a zero outcome can derive from two different processes. In our model the two processes are: macrophages had not been infected by *Leishmania* (not infected); macrophages had been infected by *Leishmania*, but have the potential to clear the infection (infected). For the not infected, the only possible outcome is zero. For the infected ones, it is a count process with integer values >=0. The two parts of the zero-inflated model form a binary model: a logit model, to determine to which of the two processes the zero outcome is associated with; a count-negative binomial model, to analyze the count outcome. The expected counts derive from the combination of the two processes (see Supplementary Information for supplementary references).

### Determination of M1 and M2 cytokines and NO production

All the cytokines were determined by ELISA kits: IL-12p40, IL-10 (Biolegend, USA), IL-1β, IL-6, TNFα (Thermo Fisher, USA), according to manufacturer’s instructions. Only for IL-10 and IL-1β quantification, the cells were pre-stimulated with LPS, 1 μg/ml for 12h, before bacterial stimulation and/or *Leishmania* infection. Simultaneous determination of nitrate and nitrite concentrations induced by the bacteria was done by Vanadium assay with the reduction of nitrate to nitrites by Vanadium (III) combined with detection by the acidic Griess reaction (Sigma-Aldrich, USA), as reported in Miranda et al., 2001. In brief, saturated solutions of Vanadium (III) chloride (VCl_3_) were prepared in 1M HCl. Culture supernatants, collected at 24h and 48h of co-infection, were mixed with the same volume of VCl_3_ and reacted with an equal volume of the Griess reagents. The absorbance at 540 nm was measured using a plate reader following the incubation. The analyses of the data were performed applying the ES approach, as deeply described in the previous section “Phagocytosis Assay and Electron microscopy”.

### Arginase and iNOS expression by real time quantitative PCR

To evaluate arginase expression, cells were pre-treated with IL-4 (200 U/ml) (R&D Systems, Minneapolis, MN) for 12h at 37°C and then stimulated and infected as above, while for iNOS (inducible nitric oxide synthase) expression no priming was performed. After 24h of infection for iNOS and 48h for arginase, the cells were collected. Total RNA was extracted using the ReliaPrep™ RNA Tissue (Promega, Madison, WI, USA) following manufacturer’s instructions. cDNAs were synthesized from RNA using the LunaScript™ RT SuperMix Kit from New England BioLabs (NEB, USA) according to the manufacturer’s instructions. Quantitative real-time PCR (qPCR) was performed using a BioRad CFX Real-Time PCR Detection System (Bio-Rad, USA) using *β-Actin* and *cyclophilin* genes for normalization (Supplementary Table 5), according to standard PCR conditions and software analysis.

### ROS determination

Intracellular reactive oxygen species (ROS) were measured by a fluorometric assay using 2’,7’-dichlorodihydrofluorescein diacetate (H_2_DCF-DA) as probe. In brief, macrophages were seeded (35,000/well) in a final volume of 200 μl/well in 96-well microplates and allowed to adhere 24h at 37°C. After an overnight incubation, the supernatants were discarded, and the cells were washed with PBS. H_2_DCF-DA was added to macrophages, and incubated 1h at 37°C. Subsequently, the cells were washed and stimulated with the two strains of bacteria and then infected with Leishmania, as reported above. The cells were incubated, protected from light, for approximately 14h. Half an hour before ending of incubation, a group of cells was treated with 1mM H2O2 and the fluorescence was measured at 485 nm (Ex) / 535 nm (Em). The analyses of the data were performed applying the ES approach, as described in the previous section “Phagocytosis Assay and Electron microscopy”.

### Cell surface markers analysis by flow cytometry

Expression of CD40, CD80, CD86 and MHC-II was evaluated by Cytofluorimetric Analysis using FACSCanto II cytometer (Becton Dickinson, Franklin Lakes, NJ). For the evaluation of co-stimulatory molecules, the cells were harvested 24h post infection with bacteria and Leishmania as indicated previously, washed with PBS and stained with appropriate dilutions of the following fluorochrome-conjugated antibodies: CD40-PE, CD80-Alexa Fluor488 and CD86-PE/Cy7 (Biolegend San Diego, CA) for 15 min at 37°C. The cells were washed, resuspended in the FACS washing buffer and finally analyzed. LPS treatment (0.3 μg/ml) was used as control. For the evaluation of MHC-II molecules, the cells were pre-stimulated with INF-γ (1 ng/ml) 12h before the infection. After 48h of co-infection the cells were harvested and processed as above using an anti-MHC-II FITC-conjugated antibody (Biolegend San Diego, CA). Treatment with PMA (50 ng/ml) was used as positive control. FACS data were analyzed with FlowJo software (TreeStar, Ashland, Ore). The differences in the proportions of macrophages presenting CD40, CD80, CD86 or MHC-II between the different treatment groups were analysed by a generalized linear mixed model (GLMM) with two random effects: the biological and the technical replicates, and the treatment with LPS or PMA as the reference effect, using poisson family as log link function. e^(intercept+estimated coeff)^ can be interpreted as the proportion of cells in the different treatment groups (see Supplementary Information for supplementary references).

## Acknowledgments

The authors thank Prof. Donatella Taramelli, Prof. Elena Crotti, Prof. Guido Favia, Dr. Stefania Orsini and Dr. Emanuela Clementi for their suggestions.

## Author contributions

C.B. and S.E. conceived and coordinated the study. I.V.B, I.A. and S.E. designed and performed the majority of the experiments. Y.C., N.B., M.G., L.G. contributed to cell culture and *Leishmania* maintenance. L.S. performed TEM analysis. R.N., M.P. contributed to phagocytosis assays and flow cytometry analyses. L.G. contributed to the design of the assay on Leishmania killing. P.G., V.T. analyzed all the data. I.V.B, C.B. and S.E. wrote the paper with input from all of the authors.

