## Supplementary Information for "Boosting innate immunity: *Asaia* bacteria expressing a protein from *Wolbachia* determine macrophage activation and killing of *Leishmania*"

PNAS
www.pnas.org

**Supplementary information**

**Boosting innate immunity: *Asaia* bacteria expressing a protein from *Wolbachia* determine macrophage activation and killing of *Leishmania***

Ilaria Varotto-Boccazzi[1#], Sara Epis[1*#], Irene Arnoldi[1,2], Yolanda Corbett[1], Paolo Gabrieli[1], Moira Paroni[3], Riccardo Nodari[1], Nicoletta Basilico[4], Luciano Sacchi[2], Marina Gramiccia[5], Luigi Gradoni[5], Vito Tranquillo[6], Claudio Bandi[1]

This PDF file includes:

Supplementary Figures 1-3
Supplementary Table 1-5
Supplementary Text
Supplementary References

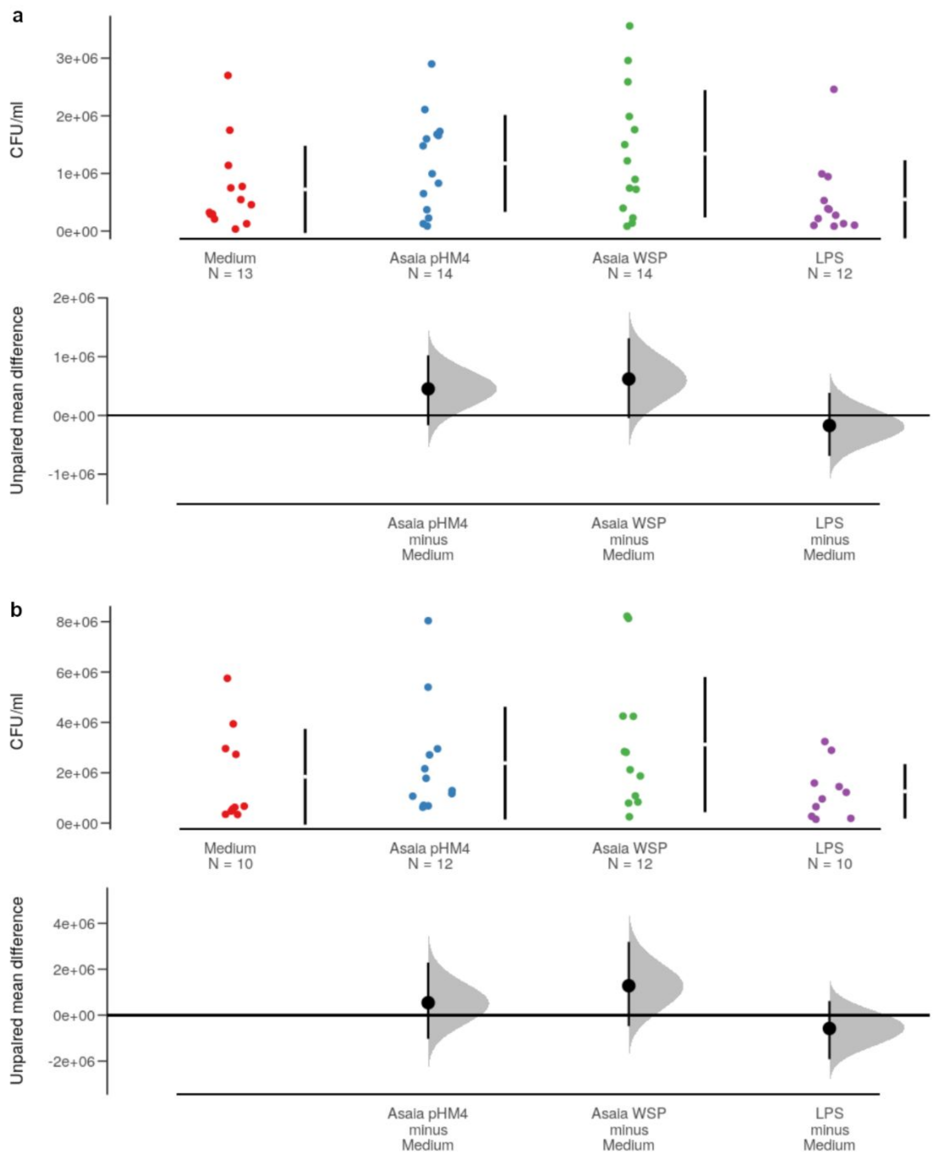

**Supplementary Fig. 1. *Staphylococcus epidermidis* uptake by macrophages pre-stimulated with *Asaia* bacteria.** Murine macrophages were pre-stimulated with *Asaia*$^{WSP}$, *Asaia*$^{pHM4}$, LPS or unstimulated (Medium) and then incubated with *S. epidermidis* bacteria for 1h (a) or 2h (b). In the top panels of the estimation plot the numbers of *S. epidermidis* bacteria internalized within macrophages are reported as CFU/ml. The vertical error bars denote mean and standard deviation of the observed data. In the bottom panels the unpaired mean differences between treatment groups compared to *Asaia*$^{pHM4}$ are reported. The shaded distribution derives from the application of the resampling algorithm, the large black circle represents the average difference between groups, the error bars on the large black circle indicate the 95% confidence interval for the calculated difference. Estimation Statistics (ES) approach and bootstrapped Welch two-sample t-test were applied.

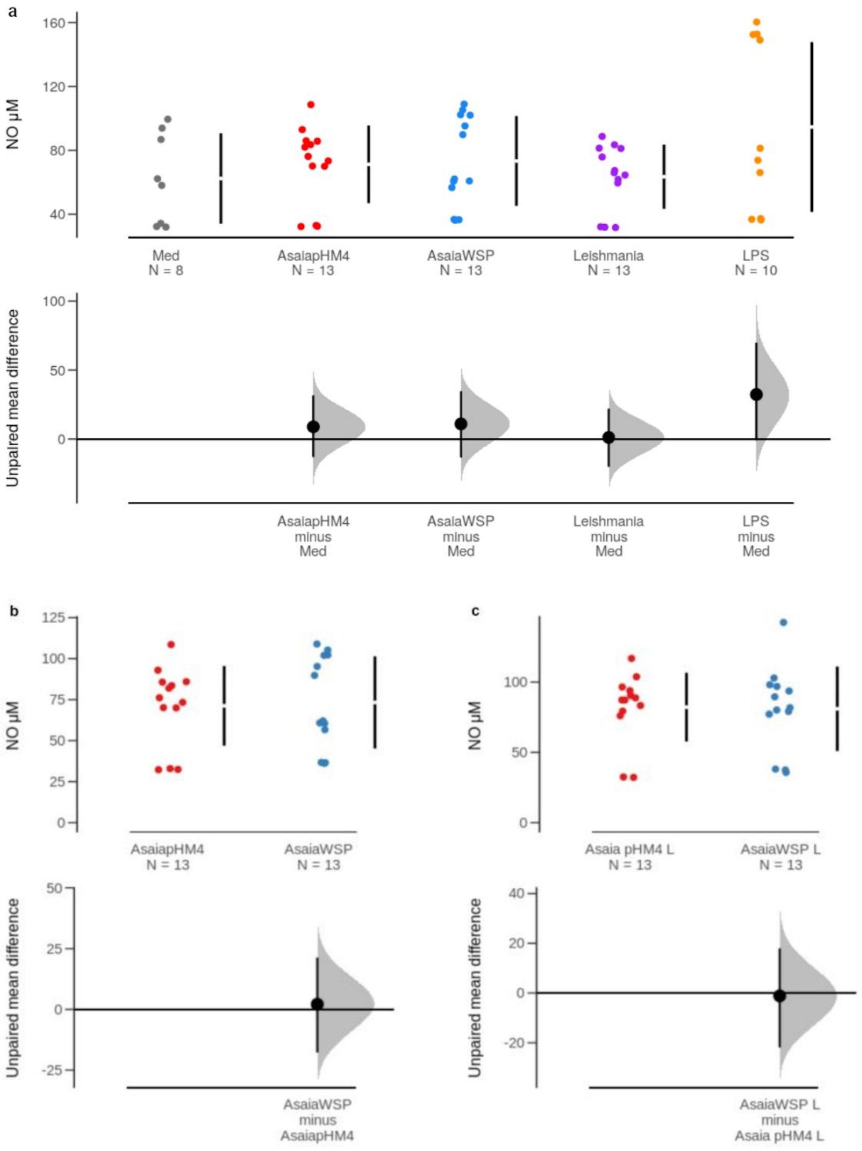

**Supplementary Fig. 2. Nitrites production by macrophages incubated with *Asaia*$^{WSP}$ and *Asaia*$^{pHM4}$ after 24h of infection.** a) Observed data (Log µM) of nitrites levels produced by macrophages stimulated with *Asaia* bacteria, *Leishmania*, LPS or untreated (Med) are reported at the top. The graphs below show the unpaired mean comparisons of the nitrite levels between the different groups and the control (untreated macrophages). b) At the top observed data (µM) of nitrites production by macrophages incubated with *Asaia*$^{WSP}$ or *Asaia*$^{pHM4}$ in absence of *Leishmania* are shown. At the bottom panel, the unpaired mean comparisons between *Asaia*$^{WSP}$ and *Asaia*$^{pHM4}$, as control, are reported. c) Observed data (µM) of nitrites production by macrophages incubated with *Asaia*$^{WSP}$ or *Asaia*$^{pHM4}$ in presence of *Leishmania* and unpaired mean comparisons are shown at the top and at the bottom, respectively. The vertical error bars denote the 95% confidence intervals of the observed data. ES approach and bootstrapped Welch two-sample t-test were applied. Data are representative of three independent experiments.

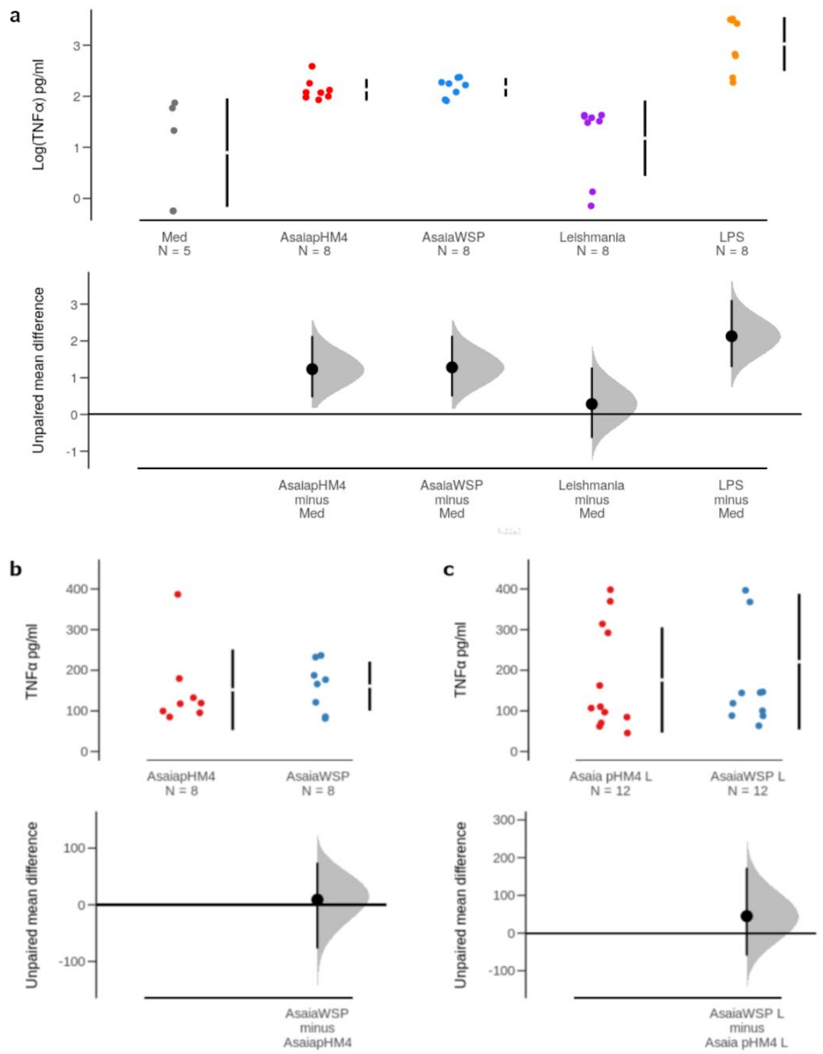

**Supplementary Fig. 3. TNFα cytokine released by macrophages incubated with *Asaia* bacteria and *Leishmania* after 48h of infection.** a) Observed data (Log pg/ml) of TNFα levels produced by macrophages exposed to *Asaia* bacteria, *Leishmania*, stimulated with LPS or untreated (Med) are reported at the top. The graphs below show the unpaired mean comparisons of the TNFα levels between the different groups and the control (Med). b) At the top observed data (pg/ml) of TNFα production by macrophages incubated with *Asaia*$^{WSP}$ or *Asaia*$^{pHM4}$ in absence of *Leishmania* are showed. At the bottom panel the unpaired mean comparisons between *Asaia*$^{WSP}$ and *Asaia*$^{pHM4}$, as control, are reported. c) Observed data (pg/ml) of TNFα production by macrophages incubated with *Asaia*$^{WSP}$ or *Asaia*$^{pHM4}$ in presence of *Leishmania* and unpaired mean comparisons are shown. The vertical error bars denote mean and standard deviation of the observed data. ES approach and bootstrapped Welch two-sample t-test were applied. Data are representative of three independent experiments.

**Supplementary Table 1a**: ***Asaia*** **phagocytosis: numbers (mean and SD) of bacteria internalized within macrophages expressed as CFU/ml and relative statistics**

| | | Treatments (CFU/ml) | | |
|---|---|---|---|---|
| | Time | Asaia pHM4 | Asaia WSP | Asaia pHM4 +LPS |
| Mean | 1h | 3,25E+05 | 7,66E+05 | 1,39E+06 |
| Std. Deviation | 1h | 1,18E+05 | 5,22E+05 | 1,06E+06 |
| Mean | 2h | 4,85E+05 | 8,09E+05 | 8,20E+05 |
| Std. Deviation | 2h | 3,09E+05 | 5,17E+05 | 5,01E+05 |
| Source of Variation | % of total variation | P value | P value summary | Significant? |
| Interaction | 2,036 | 0,4351 | ns | No |
| Bacteria | 19,56 | 0,0007 | *** | Yes |
| hours | 0,06108 | 0,8228 | ns | No |

**Supplementary Table 1b**: ***Asaia*** **survival: number (Log CFU/ml) of phagocyted bacteria in 2h and survived until 24h and relative statistics**

| | Treatments (CFU/ml) | | |
|---|---|---|---|
| | Asaia pHM4 | Asaia WSP | Asaia pHM4+ LPS |
| Mean | $2.03x10^4$ | $1.43x10^4$ | $1.07x10^4$ |
| Std. Deviation | $3.15x10^4$ | $8.9x10^3$ | $8.24x10^3$ |
| GLOBAL TEST P VALUE: 0.6279 | | | |

**Supplementary Table 2a: infection rate (mean and SD) after 48h and 24h of incubation**

| | Infection rate (percentage of infected macrophages) | | | |
|---|---|---|---|---|
| | *48h co-incubation* | | *24h co-incubation* | |
| | Mean | Standard deviation (SD) | Mean | Standard deviation (SD) |
| Leishmania | 48,0 | 16,2 | 52,4 | 16,0 |
| AsaiapHM4 | 39,6 | 10,7 | 48,4 | 13,3 |
| AsaiaWSP | 34 | 7,159 | 47,4 | 15,8 |
| Amphotericin B | 20,7 | 3,09 | - | - |

**Supplementary Table 2b: number of amastigotes (mean and SD) per infected macrophage after 48h and 24h of incubation**

| | N° Leishmania amastigotes per infected macrophage | | | |
|---|---|---|---|---|
| | *48h co-incubation* | | *24h co-incubation* | |
| | Mean | Standard deviation (SD) | Mean | Standard deviation (SD) |
| Leishmania | 2,77 | 0,95 | 2,80 | 0,59 |
| AsaiapHM4 | 2,56 | 0,87 | 2,64 | 0,99 |
| AsaiaWSP | 1,88 | 0,39 | 2,42 | 0,94 |
| Amphotericin B | 1,61 | 0,21 | - | - |

**Supplementary Table 2c: coefficient and corresponding mean number of amastigotes per macrophage after 48h and 24h of incubation**

| | N. of *Leishmania* amastigotes per macrophage | | | | | | | |
|---|---|---|---|---|---|---|---|---|
| | *48h co-incubation* | | | | *24h co-incubation* | | | |
| *Predictors* | *Coefficient (log-mean)* | *95%CI* | *Mean* | *p-value* | *Coefficient (log-mean)* | *95%CI* | *Mean* | *p-value* |
| **Count Model (log-Mean)** | | | | | | | | |
| Leishmania (Intercept) | 0.48 | 0.27 – 0.69 | 1.62 | | 1.06 | 0.52 – 0.81 | 1.93 | <0.001 |
| group [AsaiapHM4] | -0.11 | -0.29 – 0.08 | 1.45 | 0.265 | -0.04 | -0.20 – 0.12 | 1.86 | 0.601 |
| group [AsaiaWSP] | -0.88 | -1.10 – -0.66 | 0.67 | <0.001 | -0.14 | -0.31 – 0.02 | 1.68 | 0.080 |
| group [Amphotericin B] | -1.27 | -1.59 – -0.96 | 0.45 | <0.001 | - | - | - | - |
| **Zero-Inflated Model (log-odds)** | | | | | | | | |
| Leishmania (Intercept) | -2.12 | -3.90 – -0.33 | | | -1.32 | -1.87 – -0.78 | | <0.001 |
| group [AsaiapHM4] | 0.90 | -0.28 – 2.08 | | 0.135 | 0.28 | -0.18 – 0.75 | | 0.230 |
| group [AsaiaWSP] | -0.79 | -4.29 – 2.72 | | 0.660 | 0.22 | -0.26 – 0.70 | | 0.365 |
| group [Amphotericin B] | 1.12 | -0.28 – 2.53 | | 0.118 | - | - | | - |

### Supplementary Table 3a: iNOS relative expression

| | iNOS relative expression | |
|---|---|---|
| | **Mean** | **Standard deviation** |
| AsaiaWSP | 1.35 | 0.56 |
| AsaiapHM4 | 0.2 | 0.21 |
| Asaia WSPL | 0.6 | 0.41 |
| Asaia Phm4L | 0.14 | 0.01 |
| Leishmania | 0.00219 | 0.00326 |
| Medium | 0.05 | 0.06 |
| LPS | 3.04 | 1.58 |

### Supplementary Table 3b: IL10 production

| | IL-10 | |
|---|---|---|
| | **Mean (pg/ml)** | **Standard deviation** |
| AsaiaWSP | 565.81 | 222.3 |
| AsaiapHM4 | 692.05 | 75.3 |
| Asaia WSPL | 529.74 | 115.68 |
| Asaia Phm4L | 516.36 | 32.84 |
| Leishmania | 363.13 | 309.61 |
| Medium | 2.72 | 0.59 |
| LPS | 491.36 | 160.44 |

### Supplementary Table 3c: arginase relative expression

| | Arginase relative expression | |
|---|---|---|
| | **Mean** | **Standard deviation** |
| *Asaia*WSP | 1.02 | 0.14 |
| *Asaia*pHM4 | 0.54 | 0.62 |
| *Asaia* WSPL | 0.37 | 0.07 |
| *Asaia* pHM4 L | 0.76 | 0.16 |
| Leishmania | 0.28 | 0.04 |
| Medium | 0.88 | 0.31 |
| IL-4 | 25.75 | 5.45 |

#### Supplementary Table 4a: random effects used for the flow cytometry analyses

| Random Effects | CD40+ cells | CD80+ cells | CD86+ cells | MHC+ cells |
|---|---|---|---|---|
| σ2 | 0.24 | 0.51 | 0.42 | 0.05 |
| τ00 | 0.24 well:plate | 0.51 well:plate | 0.42 well:plate | 0.05 well:plate |
| | 0.17 plate | 0.82 plate | 0.17 plate | 0.11 plate |
| ICC | 0.41 | 0.62 | 0.29 | 0.67 |
| N | 9 well | 9 well | 9 well | 8 well |
| | 4 plate | 4 plate | 4 plate | 3 plate |
| Observations | 24 | 24 | 24 | 18 |

#### Supplementary Table 4b: estimated coefficients and p value of flow cytometry analyses

| | cellCD40+ | | cellCD80+ | | cellCD86+ | | cellMHC+ | |
|---|---|---|---|---|---|---|---|---|
| Predictors | Estimate | p | Estimate | p | Estimate | p | Estimate | p |
| (Intercept) | -0.24 | 0.421 | -1.10 | | -0.52 | | -1.23 | |
| Controllo positivo | Reference | | Reference | | Reference | | Reference | |
| Medium | -2.25 | <0.001 | -1.27 | 0.008 | -2.60 | <0.001 | -1.92 | <0.001 |
| Leishmania | -1.46 | <0.001 | -1.33 | 0.003 | -1.36 | 0.001 | -0.18 | 0.320 |
| AsaiapHM4 +L | -0.31 | 0.323 | -0.64 | 0.158 | -0.54 | 0.188 | 0.11 | 0.545 |
| Asaia WSP +L | -0.12 | 0.688 | -0.40 | 0.377 | -0.39 | 0.344 | 0.39 | 0.027 |

**Supplementary Table 5: list of primer pairs used in this study**

| AMPLIFIED GENES | FORWARD SEQUENCES 5'-3' | REVERSE SEQUENCES 5'-3' | REFERENCES |
|---|---|---|---|
| **iNOS** | CACCTTGGAGTTCACCCAGT | ACCACTCGTACTTGGGATGC | Tang et al., 2017 |
| **Arg I** | ACAGAGAAGGTCTCTACATCAC | CGAAGCAAGCCAAGGTTAAAGC | |
| **B-Actin** | CTCTGGCTCCTAGCACCATGAAGA | GTAAAACGCAGCTCAGTAACAGTCCG | Fallik et al., 2017 |
| **cyclophilin** | GTGACTTTACACGCCATAATG | ACAAGATGCCAGGACCTGTAT | Theret et al., 2017 |

**Supplementary Text: rationale for the selection of murine cytokines and co-stimulatory molecules**

IL-6: a protective effect by IL-6 was shown against the visceral *L. donovani* parasite (Stager et al., 2006) and a suppressive mechanism, against *L. major* infection, was highlighted in IL-6 deficient mice (Moskowitz et al., 1997). Although IL-6 is generally regarded as a Th2 cytokine, this immune mediator exerts a protective role in some forms of leishmaniases (Stager et al., 2006) and its classification into the Th1/Th2 paradigm appears problematic (Diehl & Rincón, 2002).

IL12: it is a key cytokine in the macrophage activation *in sensu* M1 being involved in the induction of nitric oxide production and expression of the enzyme iNOS; during *Leishmania* infection it is required for the maintenance of the resistance (Park et al., 2002).

TNFα and IL1-β: during leishmaniasis, TNFα is an important factor involved in the macrophage activation, killing of intracellular parasites and nitrites production; in fact, mice lacking TNFα showed fatal leishmaniasis after *L. major* infection (Wilhelm et al., 2001). The effect of TNFα is increased in presence of IL1-β, a potent inflammatory cytokine involved in the balance between inflammation and immunity which strongly affects the outcome of leishmaniasis (Dayakar et al., 2019). During *L. major* infection the production of IL1-β is downregulated and a treatment with this cytokine induces a slowing down of the disease progression in BALB/c mouse, contrary to what happens during *L. amazoniensis* infection (Ji et al., 2003).

CD40, CD80, CD86: the expression of CD40 and CD80 co-stimulatory molecules during infection with *L. major* is associated with macrophage activation and production of IL-12, which is essential for the clearance of the disease (Tuladhar et al., 2011; Barhoumi et al.,

2019). As for CD86, its role during *Leishmania* infection is critical: this molecule has been reported to participate both in defence mechanisms (Elloso & Scott 1999) as well as in the facilitation of the infection, through the induction of Th2 cytokines (Brown et al., 1996).
